# PPARγ downregulation in colonic CD8+ T cells results in epithelial barrier disruption in people with HIV on antiretroviral therapy

**DOI:** 10.1101/2024.10.02.616363

**Authors:** Upasana Das Adhikari, Leah M. Froehle, Alexandra N. Pipkin, Heeva Baharlou, Alice H. Linder, Palak Shah, Amanda Hussey, Qiming Zhang, Sarah Nyquist, Saleh Khwaled, Fangtao Chi, Swagata Goswami, Thomas J. Dieffenbach, Benjamin J. Read, Byungji Kim, Darrell Irvine, Osaretin Asowata, Mark Ladinsky, Pamela Bjorkman, Fusi Madela, Shakeel Khader, Alex Shalek, Musie Ghebremichael, Hendrik Kloverpris, Alison E Ringel, Ömer H. Yilmaz, Douglas S. Kwon

## Abstract

A hallmark of HIV infection is disruption of intestinal barrier integrity that persists in people with HIV (PWH) despite treatment with antiretroviral therapy (ART). This disruption is central to HIV disease progression, yet the causes remain incompletely understood. We report a novel mechanism by which immunometabolic defects in colon resident CD8+ T cells in PWH lead to intestinal epithelial apoptosis and disruption of intestinal barrier integrity. We show that in PWH, these cells downregulate the lipid sensor peroxisome proliferator-activated receptor-γ (PPARγ), which results in reduced intracellular lipid droplets, impaired fatty acid oxidation, and acquisition of lipids by CD8+ T cells from intestinal epithelial cells, which then contributes to epithelial cell death. Our findings indicate that HIV-associated immunometabolic dysregulation of colon CD8+ T cell leads to loss of intestinal epithelial homeostasis. These results identify potential new strategies to reduce comorbidities in PWH and other disorders with disrupted intestinal barrier integrity.

## INTRODUCTION

Intestinal barrier breakdown is a hallmark of HIV infection which persists even after long-term treatment with suppressive antiretroviral therapy (ART)^1,2^. Recent evidence points to causality, where intestinal barrier breakdown leads to the translocation of luminal microbial products into the circulation, triggering chronic systemic immune activation^3,4,5^ and significantly contributing to the development of HIV-associated non-communicable diseases (NCDs), such as diabetes, obesity, cardiovascular diseases, and stroke^6,7^. Although ART has significantly improved the lives of people with HIV (PWH), the heightened burden of NCDs continues to increase morbidity and mortality in this population^8,9^. Similar observations of chronic inflammation and loss of intestinal barrier integrity have been made in individuals with inflammatory bowel disease (IBD) and other intestinal inflammatory diseases^4,10,11^ As such intestinal barrier dysfunction is likely a key upstream driver of the diverse co-morbidities observed in PWH on ART. Despite this, the mechanisms governing intestinal barrier disruption in this population remain poorly understood.

The maintenance of intestinal barrier integrity relies on the regeneration of intestinal epithelial cells, which undergo continuous proliferation and differentiation. This process is influenced by external signals from both epithelial and non-epithelial cells, particularly stromal cells and tissue resident immune cells^12–15^. Studies suggest T cells play an important role in maintaining tissue homeostasis, including surveying the epithelium, and eliminating intestinal epithelial stem cells with aberrant behavior through interactions with MHC class I^16,17^. However, the potential role of resident T cells in the maintenance or disruption of the intestinal epithelium in PWH is incompletely understood.

In this study, we identify a novel mechanism by which colon tissue resident memory (TRM) CD8+ T cells contribute to impaired intestinal barrier integrity in PWH on ART. We obtained endoscopic biopsies from human subjects and show increased colon epithelial cell apoptosis *in vivo* and in patient-derived organoids from PWH. Our results show that colon TRM CD8+ T cells in PWH on ART downregulate expression of the lipid sensor peroxisome proliferator-activated receptor-γ (PPARγ), which leads to impaired cellular lipid metabolism. This metabolic dysregulation of colon resident CD8+ T cells contributes to intestinal epithelial cell apoptosis through a non-canonical interaction between T cells and epithelial cells. We further demonstrate the importance of PPARs in this process using a murine genetic knockout model and show that PPAR-dependent metabolic dysregulation in colon TRM CD8+ T cells results in epithelial cell apoptosis and impaired barrier integrity.

## RESULTS

### Intestinal barrier disruption in people with HIV (PWH) on antiretroviral therapy (ART) is mediated by colon resident immune cells

We observed elevated plasma intestinal fatty acid binding protein (I-FABP), a biomarker of intestinal epithelial damage, in PWH who were ART naïve or treated compared to HIV-uninfected individuals (**Figure 1A**). To examine colon epithelial damage in those with HIV, we stained colonic tissue biopsies for the apoptosis marker cleaved caspase 3 (CC3) and found increased cell death in the epithelium of both ART naïve and ART treated PWH compared to those without HIV infection (**Figures 1B, 1C**). Despite these findings, the frequency of intestinal stem cells and proliferative cells in colon crypts remained unchanged with HIV infection (**Figure S1A-D**). We also found no evidence of aberrant tight junctions in PWH on ART using electron microscopy of the colon epithelium (**Figure S1E**). To further assess the mechanism of epithelial apoptosis in PWH, we established a patient-derived colon organoid (“colonoid”) model using endoscopic colon tissue biopsies (**Figure 1D**). Primary colonoids derived directly *ex vivo* from PWH on ART exhibited greater epithelial apoptosis than those from HIV-uninfected individuals (**Figures 1E**). When these colonoids were mechanically dissociated to single cells, epithelial stem cells in the primary colonoids of PWH on ART had the potential to regrow into new colonoids (referred to as “secondary colonoids”) (**Figure 1F**). This capacity of the intestinal stem cells to form secondary colonoids (“clonogenicity”) was the same between PWH on ART and HIV-uninfected individuals (**Figure S1F**). Interestingly, the increase in epithelial apoptosis observed in primary colonoids derived from PWH on ART was no longer seen in their secondary colonoids (**Figures 1G and H**). Consistent with this, lactate dehydrogenase (LDH) release, a measure of cellular damage, was increased in primary colonoids from PWH on ART compared to those from HIV-uninfected individuals, but this was not observed in secondary colonoids (**Figure S1G**). These results indicate that intestinal epithelial cell apoptosis observed *in vivo* was recapitulated in primary colonoids derived from PWH on ART, but this cellular death was lost after disassociation and seeding of secondary colonoids.

**Figure 1.**
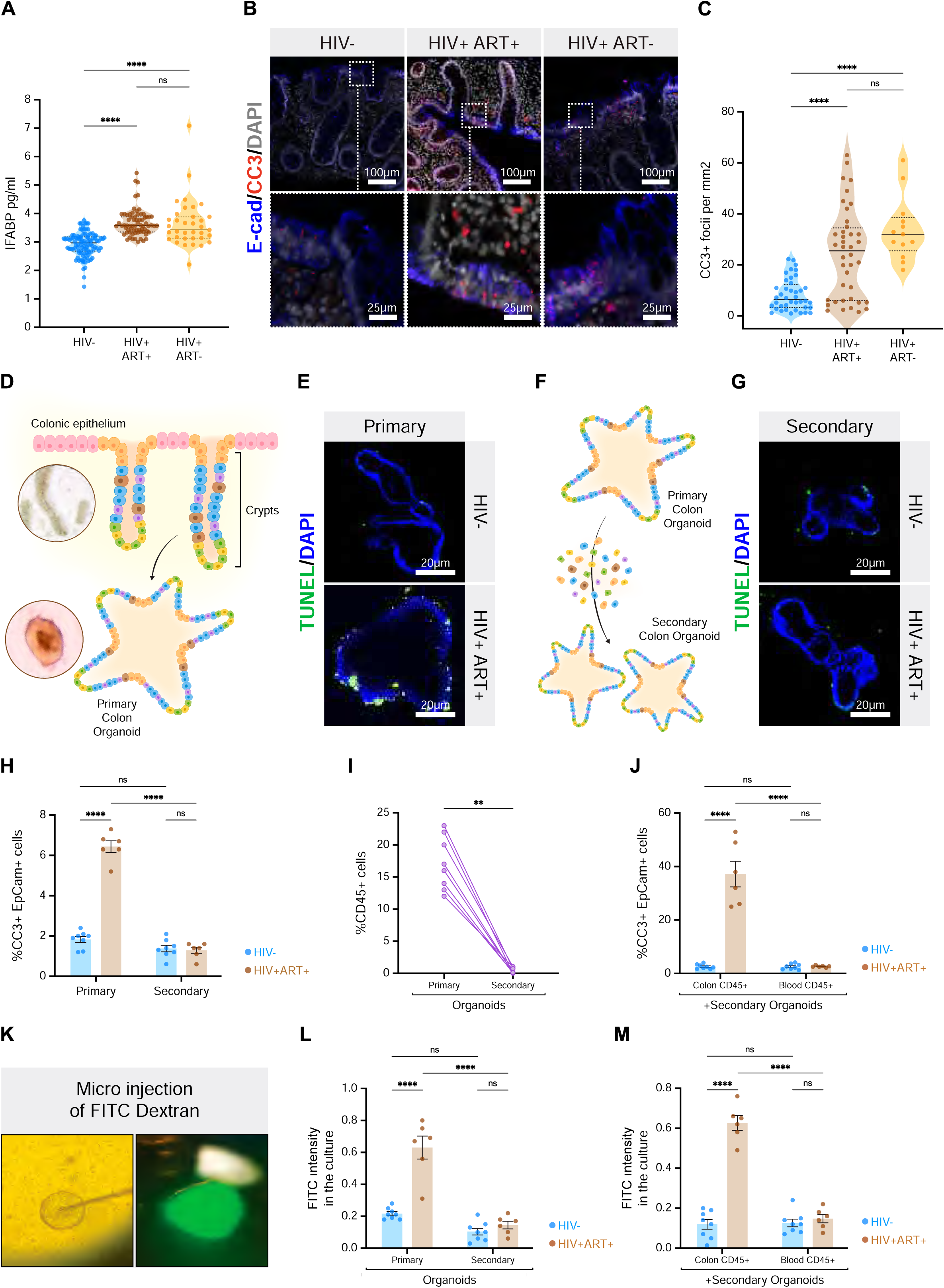
Intestinal barrier disruption in people with HIV (PWH) on antiretroviral therapy (ART) is mediated by colon resident immune cells A. IFABP concentration in plasma among PWH with ART, PWH without ART and uninfected individuals. B. Confocal microscopic images of colonic biopsy showing epithelial apoptosis (E-cadherin/Cleaved caspase 3) among PWH with ART, PWH without ART and uninfected individuals. Scale bars 100μm (top row); 25μm (bottom row) C. Quantification of confocal imaging for colonic epithelial apoptosis among PWH with ART, PWH without ART and uninfected individuals. D. Schematic showing primary colonoid establishment E. Increased apoptosis (TUNEL+) in primary colonoids of PWH on ART compared to uninfected patient derived colonoids. Scale bar 20μm F. Schematic showing secondary (post passaged) colonoid establishment G. Secondary colonoids of PWH on ART display indifferent epithelial cell death phenomenon in PWH on ART compared to uninfected. Scale bar 20μm H. Quantification of flow cytometric analysis of colonic epithelial apoptosis (CC3+ Epcam+) from the colonoid cultures. I. Presence of CD45+ immune population in primary colonoids vs secondary cultures. J. Colonic CD45+ immune cells induce epithelial apoptosis unlike blood derived immune cells from PWH. K. Microinjection of FITC dextran into the lumen of colonoids. L. Increased FITC intensity in the primary organoid cultures of PWH compared to uninfected and secondary organoids from PWH respectively. M. Increased FITC intensity in the secondary organoid culture of PWH when cocultured with patient autologous CD45+ immune cells.

To further assess differences in primary and secondary colonoids, we characterized their cellular composition by flow cytometry and identified a loss of CD45+ immune cells in secondary colonoids (**Figure 1I**). Increased epithelial apoptosis was restored when autologous colon CD45+ immune cells from the lamina propria but not peripheral blood were co-cultured with the secondary colonoids from PWH on ART (**Figure S1H, Figure 1J**). Addition of autologous colon CD45+ immune cells to secondary organoids derived from HIV-uninfected individuals did not result in epithelial apoptosis (**Figure 1J**). This was again consistent with observations made examining LDH release (**Figure S1I**). These findings indicate that colon CD45+ immune cells were necessary and sufficient to induce the epithelial apoptosis observed in colonoids derived from PWH on ART.

To determine whether colon epithelial apoptosis in PWH on ART led to a loss of barrier integrity, FITC-dextran was microinjected into the lumen of primary colonoids to measure leakage into the supernatant (**Figure 1K)**. We observed increased FITC leakage in primary colonoid cultures from PWH on ART compared to those from HIV-uninfected individuals (**Figure 1L**). FITC leakage was lost in secondary colonoids but could be recapitulated with addition of autologous colon resident CD45+ immune cells but not autologous blood derived CD45+ cells from PWH on ART (**Figure 1M**). Together, these findings indicate that colon resident CD45+ immune cells from PWH on ART mediate epithelial apoptosis and disruption of barrier integrity.

### Colon CD8+ T cells from PWH on ART induce epithelial apoptosis via non-canonical functions

To determine which specific CD45+ immune cell subset meditated epithelial apoptosis, we co-cultured secondary colonoids from PWH on ART with autologous colon CD4+ T cells, CD8+ T cells, and non-T immune cells (CD45+ CD3-). Only inclusion of CD8+ T cells recapitulated the epithelial damage observed in primary colonoids (**Figure 2A**). We then used confocal microscopy to examine determine the localization of CD8+ T cells relative to the epithelium in colon tissue samples. We found that in PWH on ART, colon CD8+ T cells were localized closer or within the epithelial layer compared to those who were HIV-uninfected (**Figure 2B, 2C, and S2A**). There was no significant difference in the frequency of total CD8+ T cells in the colon between PWH and uninfected individuals (**Figure S2B**). To better understand how CD8+ T cells were mediating this epithelial apoptosis, we cultured colonoids with autologous CD8+ T cells either separated by a transwell or together, or with T cell supernatants. This demonstrated that epithelial apoptosis was primarily dependent upon direct contact of CD8+ T cells and epithelial cells, although there may be some contribution from soluble factors (**Figure 2D**). These findings indicate that close proximity of colon CD8+ T cells to the epithelium is necessary for epithelial apoptosis in PWH on ART.

**Figure 2.**
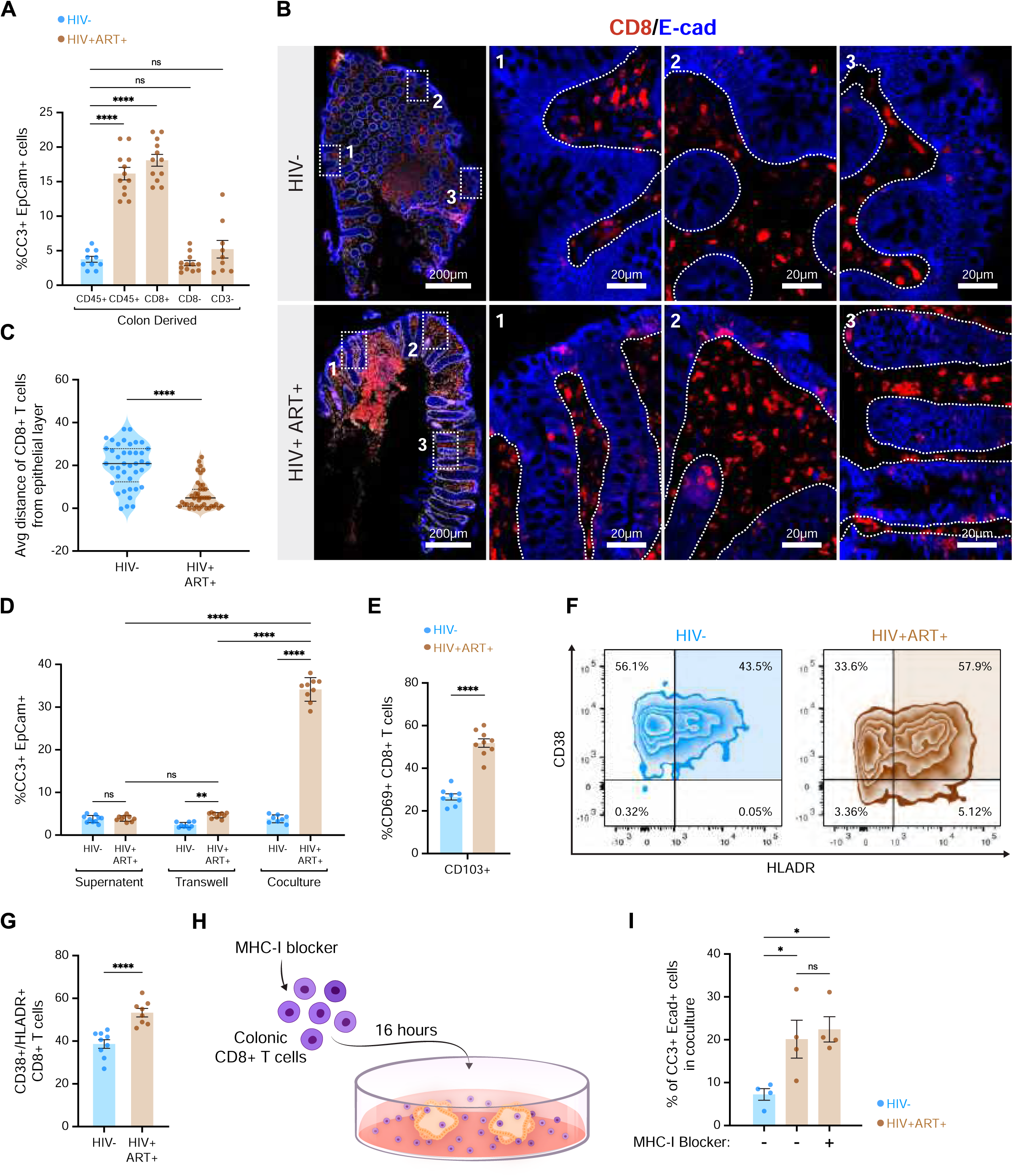
Colonic CD8+ T cells from PWH on ART trigger epithelial apoptosis through a contact-dependent, non-canonical mechanism. A. Flow cytometric quantification of the frequency of epithelial apoptosis (EpCam+ CC3+) in coculture of colonic immune subpopulations with patient autologous secondary colonoids derived from PWH on ART. B. Confocal microscopy images of colon biopsy pinches showing the localization of CD8+ T cells and the epithelial region. Scale bar 200μm; Insets 20μm C. Quantification of the average distance of CD8+ T cells from the epithelial layer in the colon between HIV-uninfected and PWH on ART individuals. D. Flowcytometric quantification of epithelial apoptosis (EpCam+ CC3+) in presence of colonic CD8+ T cells in transwell, coculture set up and supernatant of ex vivo colonic CD8+ T cell culture from PWH on ART and uninfected. E. Flowcytometric quantification of frequencies of colonic CD8+ CD69+ CD103+ population (colon TRM CD8+ T cells) in PWH on ART compared to uninfected F. Flowcytometric plot of colonic CD8+ CD38+ HLADR+ population in PWH on ART compared to uninfected G. Quantification of frequencies of colonic CD8+ CD38+ HLADR+ population in PWH on ART compared to uninfected H. Schematic showing MHC-I blocking of colonic CD8+ T cells prior to their coculture with colonoids I. Flowcytometric quantification of frequencies of epithelial apoptotic cells (Epcam+ CC3+) with MHC-I blocking in PWH on ART compared to uninfected

Since tissue resident memory (TRM) CD8+ T cells in the colon play a pivotal role in protection against infections and tumors, we analyzed memory (CD45RO+) resident (CD69+ CD103+/-) subsets and observed that CD103+ tissue resident memory (CD45RO+ CD69+ CD103+; hereafter “TRM” CD8+ T cells) cells were more frequent in PWH on ART than in HIV-uninfected individuals (**Figure 2E**). However, there was no difference in total memory CD8+ T cells (CD8+ CD45RO+) (**Figure S2C**). These TRM CD8+ T cells were also more activated (CD38+ HLA-DR+) in PWH on ART than HIV-uninfected people (**Figure 2F, 2G and S2D**). Notably, we found that colon derived immune cells from PWH on ART had higher levels of intracellular IL-15 compared to uninfected individuals (**Figure S2G**). This suggests T cells may be contributing to their activation, since IL-15 can stimulate T cell activation^18^. However, when TRM CD8+ T cells from HIV-uninfected people were activated (by stimulated with phytohemagglutinin (PHA), CD3/CD28 antibody, or IL-15) and co-cultured with secondary colonoids, these cells failed to induce epithelial apoptosis, indicating that T cell activation was not sufficient for this activity (**Figure S2E**). We additionally found that TRM CD8+ T cells from PWH had reduced effector cytokine, perforin, and granzyme secretion, (**Figure S2F-M)**, and blockade of these pathways did not prevent CD8+ T cell-mediated epithelial apoptosis in colonoids from PWH on ART (**Figure S2N**). Importantly, CD8+ T cell-mediated epithelial apoptosis operated independently of the MHC-I (**Figure 2H and 2I**). Overall, we demonstrate that that TRM CD8+ T cells from PWH on ART mediate epithelial cell apoptosis through a contact dependent non-canonical mechanism.

### Colon CD8+ T cells from PWH on ART exhibit impaired lipid metabolism

To further understand functional differences in colon TRM CD8+ T cells from PWH, we performed differential gene expression (DGE) analysis, which demonstrated downregulation of multiple lipid metabolism-associated genes and pathways in PWH on ART relative to HIV-uninfected individuals (**Figure 3A and 3B**). Similar findings were observed in DGE analysis of colon TRM CD8+ T cells from an independent South African cohort of PWH (**Figure S3A**). In both datasets, the peroxisome proliferator-activated receptor γ (*PPARγ*) was downregulated in PWH on ART. We also observed a significant reduction in the expression of genes downstream of PPARγ, such as *CD36*, which encodes a membrane protein that facilitates the uptake of long-chain fatty acids^19^; *Dgat2* (diacylglycerol O-acyltransferase 2), an enzyme crucial for triglyceride synthesis^20^; *Pnpla7*, which is thought to modulate intracellular lipid droplets (LDs)^21^; and *DGKα* (diacylglycerol kinase alpha) and *DGKγ* (diacylglycerol kinase gamma), enzymes that play a central role in LD formation by regulating intracellular diacylglycerol (DAG)^22^. Since PPARs are critical fatty acid (FA) sensors^23–25^, downregulation of these genes, along with reduced expression of other PPAR family members, highlights the broad disruption in lipid metabolism pathways in colon TRM CD8+ T cells in PWH on ART.

**Figure 3.**
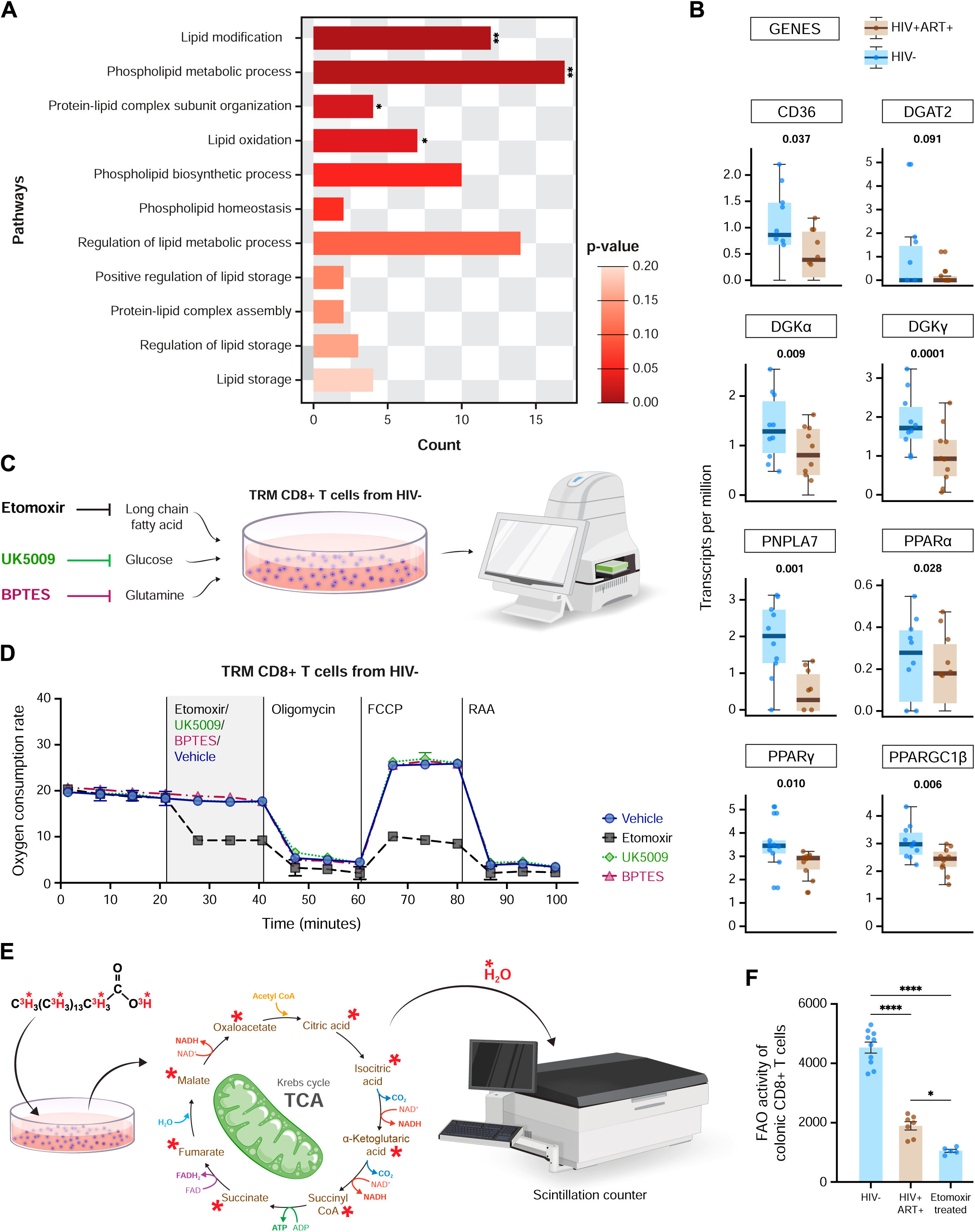
Impaired lipid metabolism in colon TRM CD8+ T cells in PWH on ART A. Pathway analysis showing gene counts that is enriched in GO terms. Benjamini-Hochberg-corrected p values: *p < 10–2, **p < 10–5. B. Log(transcripts per million) expression of PPARg downstream targets genes integral to lipid droplet biogenesis in colon TRM CD8+ T cells from HIV-uninfected individuals (blue, n = 10) and PWH on ART (brown, n = 9). C. Scheme of SeaHorse substrate oxidation assay used to assess the metabolic pathways in colon TRM CD8+ T cells from healthy, HIV-uninfected individuals. D. Oxygen consumption rate (OCR; pmol/min/cells) of sorted colon TRM CD8+ T cells from healthy, HIV-uninfected individuals was performed by SeaHorse assay to evaluate their metabolic profile. The OCR was measured during the mitochondrial stress test with the addition of oligomycin, carbonyl cyanide p-trifluoromethoxy phenylhydrazone (FCCP), and rotenone and antimycin A drugs (RAA). Results were normalized with the cell count. E. Schematic showing Fatty acid oxidation (FAO) activity by radioactive tracing of tritiated palmitic acid F. FAO activity (counts per minute [cpm]) of metabolized 3H-palmitic acid normalized to colon TRM CD8+ T cell count from HIV-uninfected and PWH on ART, accompanied by the FAO activity calculated from 4mmEtomoxir treated colon TRM CD8+ T cells from healthy, HIV-uninfected.

Due to the transcriptional dysregulation of lipid metabolism in colon TRM CD8+ T cells from PWH, we examined the metabolic requirements of these cells. We found that colon TRM CD8+ T cells in HIV-uninfected people were highly dependent upon FA utilization as demonstrated by reduced mitochondrial oxidative metabolism when incubated with etomoxir, an inhibitor of carnitine palmitoyltransferase-1A (CPT1A) which assists in shuttling of FA into the mitochondria for fatty acid oxidation (FAO)^26^ (**Figures 3C and 3D**). This was not observed in CD8+ T cells from the periphery of HIV-uninfected individuals (**Figure S3B**). The mitochondrial oxidative metabolism of colon TRM CD8+ T cells was not affected by the absence of other metabolic substrates such as glucose or glutamine (**Figures 3C and 3D**). These findings indicate that colon TRM CD8+ T cells are highly dependent upon FAO for their maintenance, unlike their peripheral counterparts.

To further investigate the role of FAO in colon TRM CD8+ T cells, we treated these cells with tritiated palmitic acid and measured the release of tritiated water as an indicator of FAO activity (**Figure 3E**). Our results demonstrated a reduction of TRM CD8+ T cell FAO in PWH on ART relative to those who were HIV-uninfected (**Figure 3F**). Treatment of CD8+ T cells from HIV-uninfected individuals with etomoxir also resulted in FAO reductions (**Figure 3F**). We next examined mitochondrial bioenergetics by measuring basal respiration and spare capacity, which are essential for meeting cellular ATP demand. Colon TRM CD8+ T cells from PWH demonstrated impaired mitochondrial oxidative metabolism, including reduced basal respiration and spare capacity compared to those from HIV-uninfected people (**Figure S3C and S3D**). Exogenous FA treatment with palmitic acid failed to restore mitochondrial oxidative metabolism in these cells (**Figure S3C and S3D**). However, colon TRM CD8+ T cells from PWH had similar FA uptake capacity when compared to HIV-uninfected individuals, as indicated by the uptake of labeled palmitic acid (BODIPY C16) (**Figure S3E and S3F**). These data suggest that colon TRM CD8+ T cells from PWH are highly dependent upon FAO and can uptake FA from their local environment but are not able to efficiently utilize exogenous FA for FAO.

### Treatment with a PPARγ agonist restores lipid homeostasis in colon CD8+ T cells from PWH on ART

Intracellular LDs act as an internal lipid reserve that supplies FAs for cellular metabolism. Given the downregulation of the lipid sensor PPARγ and its downstream targets, we explored whether LDs serve as the source of FAs for FAO in colon TRM CD8+ cells. We measured FAO in colon TRM CD8+ T cells from HIV-uninfected individuals after blocking DGAT and ATGL, two key enzymes that are involved in LD biogenesis and lipolysis, respectively (**Figure 4A**). We demonstrated that blocking these enzymes in colon TRM CD8+ T cells reduced FAO, supporting that these cells mobilize FAs from their LDs for FAO (**Figure 4B**). We also observed that using DGAT or ATGL (Atglistatin) inhibitors (**Figure S4A**) to deplete LDs also resulted in colon TRM CD8+ T cells from HIV-uninfected individuals inducing epithelial apoptosis (**Figure S4B**). This suggests that LD-mobilized FA are not only crucial for CD8+ T cell intrinsic FAO but also TRM CD8+ T cell-mediated epithelial cell apoptosis.

**Figure 4.**
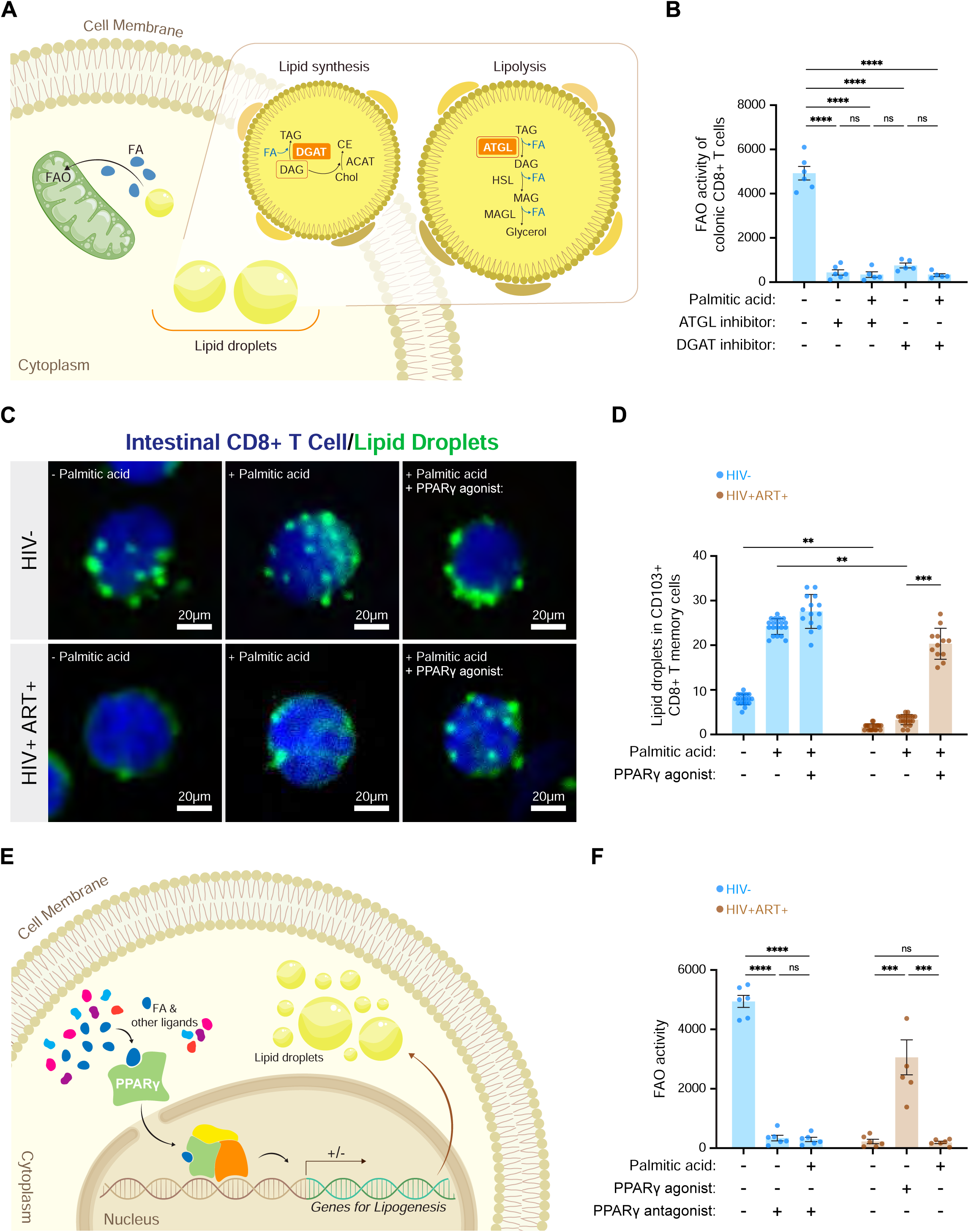
PPARγ-mediated lipid droplet repletion restores lipid homeostasis in colonic CD8+ T cells from PWH on ART A. Diagram showing lipid droplet biogenesis and lipolysis pathway, with key enzymes highlighted in orange B. Quantification of FAO activity (counts per minute [cpm]) of metabolized 3H-palmitic acid normalized to colon TRM CD8+ T cell count from healthy, HIV-uninfected individuals, upon ATGL inhibition (Atglistatin) to block lipolysis and DGAT inhibition to block lipid droplet biogenesis. C. Confocal imaging of Lipid droplets (Nile Red in green) in colon TRM CD8+ T cells from PWH on ART and uninfected; accompanied by Palmitic acid and PPARγ agonist supplementation. D. Quantification of lipid droplets in colon TRM CD8+ T cells from PWH on ART and uninfected E. Diagram showing how PPARγ ligands and fatty acids (FA) activate PPAR signaling which regulates lipid droplet dynamics. F. Quantification of FAO activity in colon TRM CD8+ T cells from healthy, HIV-uninfected individuals when treated with PPARγ antagonist (bars in blue); and colon TRM CD8+ T cells from PWH on ART when treated with PPARγ agonist (bars in brown)

We next stained colon TRM CD8+ T cells for LDs and observed LD depletion in cells from PWH on ART compared to HIV-uninfected individuals (**Figures 4C and 4D).** This depletion could not be reversed with exogenous FA (palmitic acid) (**Figures 4C and 4D**), similar to our observation that administration of exogenous FA did not enhance mitochondrial metabolism (**Figures S3C and S3D**). However, we found that treatment of these cells with the PPARγ agonist rosiglitazone replenished LDs (**Figures 4C and 4D**). This was consistent with the role of PPARγ in regulating LD biogenesis (**Figure 4E**). Treatment with rosiglitazone also increased FAO in TRM CD8+ T cells from PWH while the PPARγ antagonist GW9662 significantly reduced FAO in these cells from HIV-uninfected individual (**Figure 4F**). These findings demonstrate that colon TRM CD8+ T cells are dependent on LDs to maintain metabolic homeostasis but have depleted LDs and impaired FAO in those infected with HIV on ART. Importantly, this LD depletion and FAO defect could be reversed with the PPARγ agonist rosiglitazone.

### Colon TRM CD8+ T cell from PWH on ART mediate epithelial apoptosis through PPARγ-dependent lipid scavenging

Our findings revealed that colon TRM CD8+ T cells from PWH on ART exhibit impaired lipid metabolism. To investigate how this lipid metabolic dysregulation results in epithelial apoptosis, we performed live imaging of TRM CD8+ T cells and colonoids to observe their cellular interactions. We stained colonoid epithelial lipids with DiI, a lipophilic dye that integrates into lipid bilayers, and Cell Mask, an amphipathic dye that anchors to the plasma membrane through its hydrophilic and lipophilic components. These were co-cultured with autologous TRM CD8+ T cells labeled with Cell Trace Violet. We then measured contacts between labeled CD8+ T cells and epithelial cells and demonstrated that TRM CD8+ T cells from PWH had significantly more interactions than those from HIV-uninfected subjects (**Figure 5A, 5B**). Additionally, we also observed higher levels of transfer of DiI labeled epithelial cell lipids to TRM CD8+ T cells from PWH compared to those from HIV-uninfected individuals (**Figure 5C**). This epithelial lipid transfer to TRM CD8+ T cells was abolished in the presence of the actin polymerization inhibitor latrunculin A or the PI-3Kinase inhibitor wortmannin (**Figure 5C**). These inhibitors were previously shown to prevent capture of plasma membrane fragments from target cells by T cells and cancer cells^27,28^. Finally, treatment of colon TRM CD8+ T cells with latrunculin A or wortmannin significantly reduced epithelial apoptosis in samples derived from PWH on ART (**Figure 5D**). To further confirm the lipid scavenging of TRM CD8+ T cells, we co-cultured Cy5-PE-labeled liposomal nanoparticles mimicking plasma membranes (**Figure 5E**) and showed rapid transfer of Cy5-PE to CD8+ T cells (**Figure 5F**). These findings suggest that colon TRM CD8+ T cells from PWH on ART contact epithelial cells and acquire lipids from epithelial membranes and contribute to their apoptosis.

**Figure 5.**
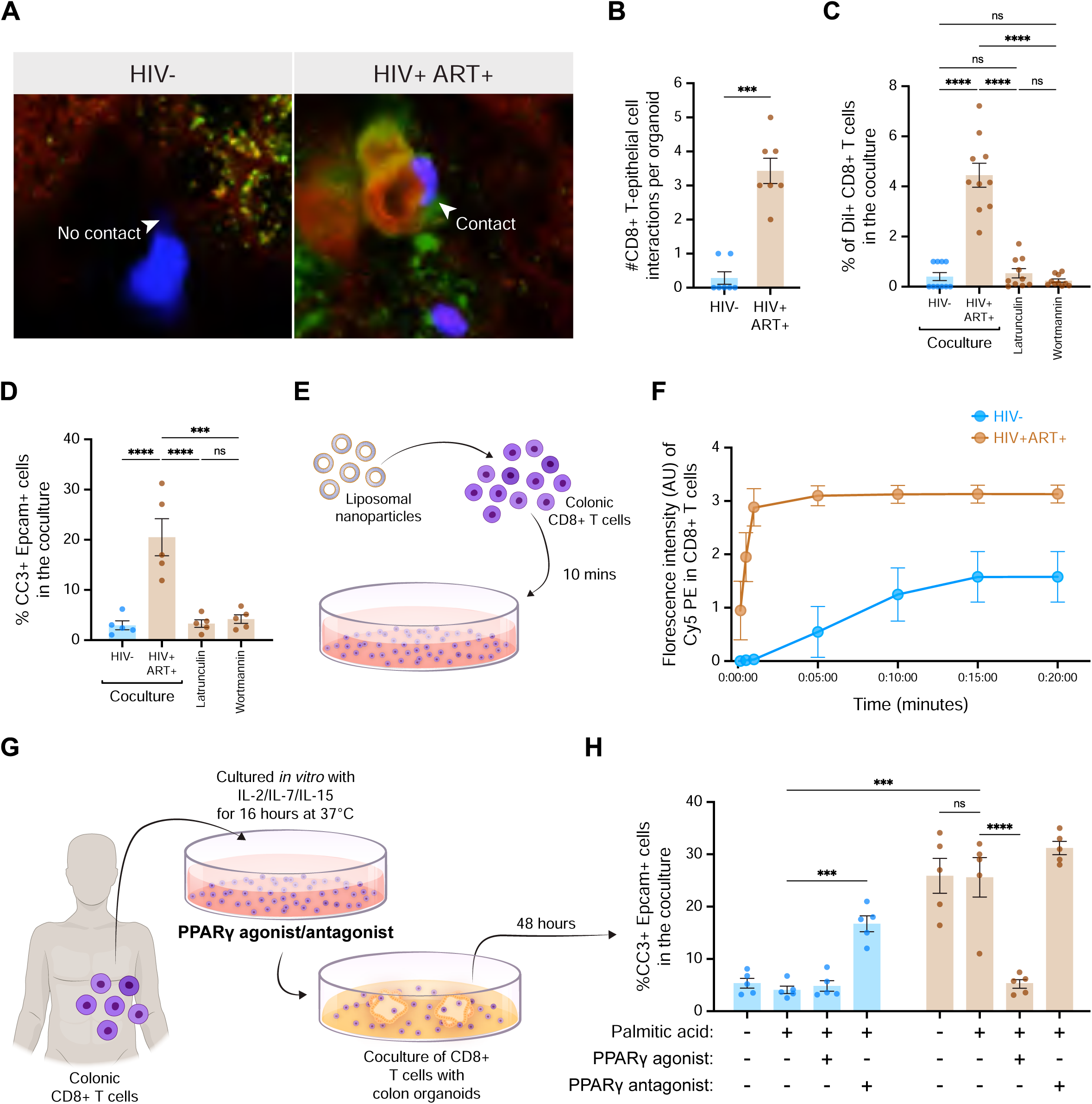
PPARγ agonist treatment to PWH derived colon TRM CD8+ T cells abrogated T cell mediated epithelial apoptosis A. Snapshots from timelapse live imaging comparing colon TRM CD8+ T cell of PWH and uninfected when cocultured with patient autologous secondary colonoids demonstrating proximity and interaction of colon TRM CD8+ T cells (blue) with epithelial cells (red) and epithelial lipid reserves (DiI). B. Quantification of the frequency of colon TRM CD8+ T interacting with the epithelial cells of the organoid. C. Quantification of the frequency of DiI+ colon TRM CD8+ T from the coculture, accompanied by coculture assays when colon TRM CD8+ T cells treated with Latrunculin (inhibitors of actin polymerization) and Wortmannin (inhibitors of PI3K) D. Flowcytometric quantification of epithelial apoptosis (Epcam+ CC3+) in coculture when colonic CD8+ T cells treated with Latrunculin (inhibitors of actin polymerization) and Wortmannin (inhibitors of PI3K) E. Schematic showing colon TRM CD8+ T cells treated by fluorophore tagged liposomal nanoparticles F. Quantification of uptake frequency of liposomal nanoparticles by colon TRM CD8+ T cells. G. Schematic showing colon TRM CD8+ T cell treated with PPARγ antagonist and agonist and then cocultured with autologous patient derived secondary colonoids. H. Flowcytometric quantification of epithelial apoptosis (Epcam+/ CC3+) in coculture; when colon TRM CD8+ T cell of PWH on ART was supplemented with PPARγ agonist, and when colon TRM CD8+ T cell of uninfected is supplemented with PPARγ antagonist.

Having shown that restoring PPARγ in colon TRM CD8+ T cells of PWH enhances their FAO activity by replenishing their intrinsic LDs (**Figure 4F**), we aimed to determine whether treatment with a PPARγ agonist could reverse epithelial apoptosis caused by PPARγ deficient colon TRM CD8+ T cells. When colon CD8+ T cells from PWH on ART were treated with the PPARγ agonist rosiglitazone (**Figure 5G**), they induced less epithelial cell death (**Figure 5H**), decreased CD8+ T cell-epithelial contacts (**Figure S5B and S5C**), and reduced DiI transfer from the co-cultured epithelial cells (**Figure S5D**). Additionally, colon TRM CD8+ T cells from HIV-uninfected individuals treated with the PPARγ antagonist GW9662 now induced epithelial apoptosis (**Figure 5H**), had increased contacts with the epithelial cells (**Figure S5E and S5F**), and increased DiI transfer from the epithelium (**Figure S5G**). These results indicate that reduced PPARγ signaling in colon TRM CD8+ T cells results in lipid scavenging from epithelial cells, which induces epithelial cell apoptosis. Importantly, this can be reversed with the PPARγ agonist rosiglitazone.

### Colon CD8+ T cell PPAR signaling is required for maintaining intestinal barrier integrity in a murine model

To provide further support that the observed downregulation of PPARs in colon TRM CD8+ T cells in PWH on ART was critical for epithelial apoptosis, we generated a mouse lacking PPARα/β/γ (isoforms that are functionally redundant in mice) in CD8+ T cells (CD8-PPAR TKO) (**Figure S6A**). Colon CD8+ T cells from the CD8-PPAR TKO mice exhibited LD depletion, consistent with a critical role for PPARs in LD biogenesis (**Figures 6A, 6B and S6B**). We also observed a significant increase in epithelial cell apoptosis in the colon of CD8-TKO compared to WT mice (**Figure 6C and 6D**). Additionally, higher levels of epithelial apoptosis were observed with mouse colonoids were co-cultures with colonic CD8+ T cells from CD8-TKO mice compared to WT mice (**Figure S6C and S6D**). We next assessed *in vivo* intestinal barrier disruption by giving mice a FITC-dextran enema and showed that CD8-TKO mice had significantly increased barrier disruption compared to WT mice (**Figure 6E**). These findings further support a model in which downregulation of PPAR in colon TRM CD8+ T cells results in intestinal epithelial cell apoptosis and intestinal barrier disruption through heterotypic cellular interactions.

**Figure 6.**
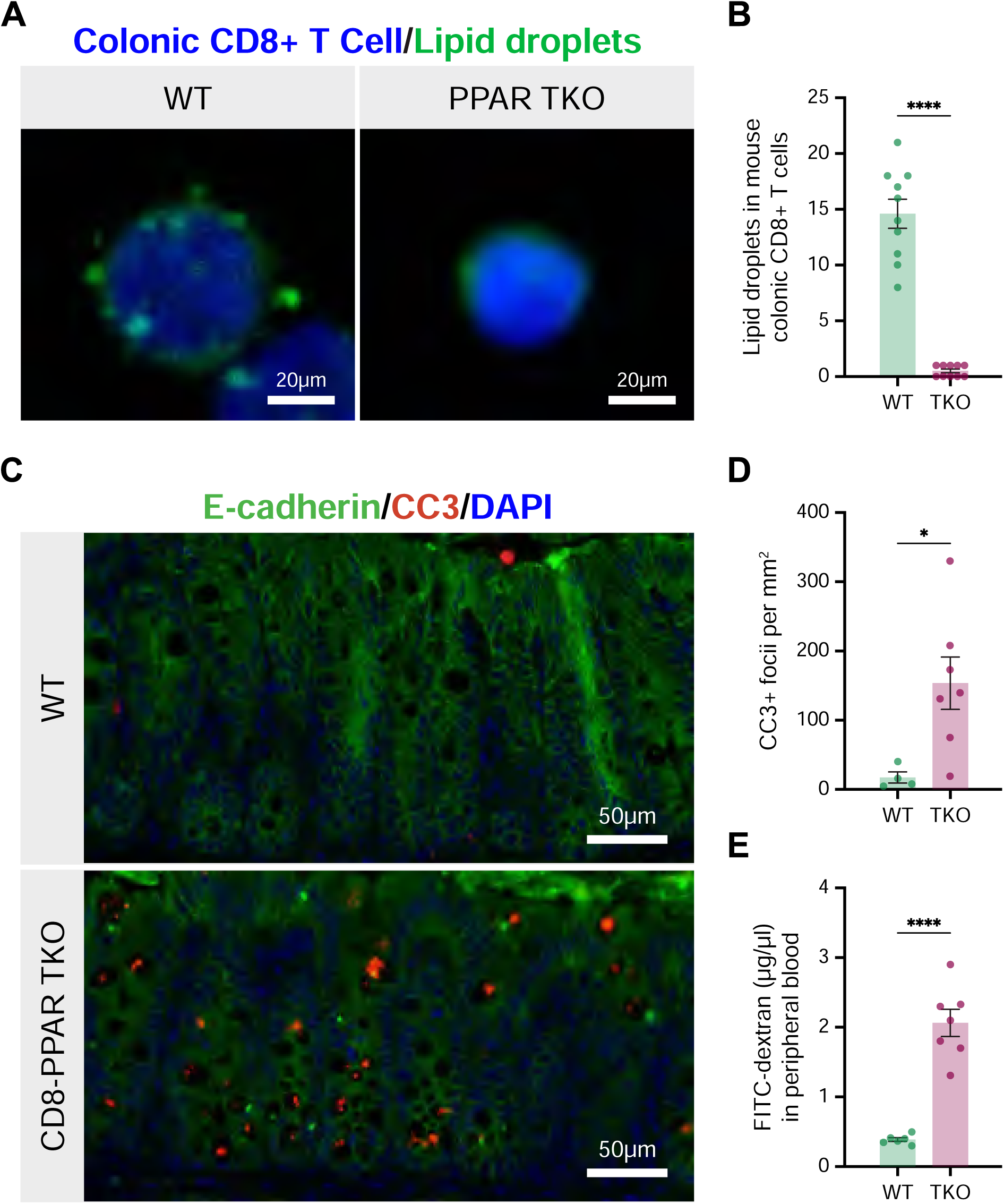
Loss of PPARγ signaling in colonic CD8+ T cells is critical for maintaining intestinal barrier integrity A. Confocal imaging of lipid droplets (green) in colonic CD8+ T cells from WT and CD8+ T cell specific PPAR TKO mice. Scale bar 20μm B. Quantification of lipid droplets in colonic CD8+ T cells from WT and PPAR TKO mice C. Confocal imaging of colon mucosa showing epithelial apoptosis (E-cadherin/CC3+) in CD8+ T cells specific PPAR TKO mice compared to WT. Scale bar 50μm D. Quantification of epithelial apoptosis between WT and CD8+ T cell specific PPAR TKO mice. E. Quantification of FITC dextran in peripheral blood of WT and CD8+ T cell specific PPAR TKO mice

## DISCUSSION

Colon resident CD8+ T cells predominantly exist as memory populations, with tissue-resident memory cells being the primary subset^29^. These cells are shaped by the local tissue microenvironment and, unlike circulating memory CD8+ T cells, are strategically positioned within the mucosa, continuously surveying the epithelium for signs of infection or abnormal cellular behavior^16^. The integrity of the intestinal barrier depends on interactions between tissue resident CD8+ T cells and the epithelium^30^. Disruption of these interactions has been implicated in various diseases, including inflammatory bowel disease (IBD), where excessive activity of colonic CD8+ T cells can compromise the intestinal epithelial barrier^10,11,32,33^. However, the impact of TRM CD8+ T cells on intestinal epithelium in PWH has not been extensively studied.

Our findings support the importance of interactions between colon TRM CD8+ T cells and epithelial cells, highlighting how metabolic dysregulation of the CD69+ CD103+ subset of resident CD8+ T cells significantly impacts the integrity of the epithelial barrier in PWH on ART. The lack of proliferation of intestinal stem cells in colon crypts during epithelial apoptosis suggests a lack of adequate compensatory mechanisms to maintain tissue homeostasis. In PWH apoptotic epithelial cells may be retained in the epithelium, leading to lack of signals to the stem cell compartment to proliferate and replace these cells, thus contributing to persistent loss of epithelial barrier integrity. Although chronic T cell activation is associated with T cell-mediated loss of epithelial integrity in cancer and IBD, this has been reported to occur via epithelial tight junction remodeling or cytokine mediated epithelial damage^31–33^. However, our findings suggest that epithelial damage in PWH requires contact with CD8+ T cells, and that epithelial death may involve non-canonical CD8+ T cell functions, suggesting potential novel mechanisms of colon TRM CD8+ T cell dysfunction during HIV infection.

Immunometabolic properties of CD8+ T cells significantly contribute to maintenance of intestinal homeostasis in PWH^34^. In the skin, it has been described that tissue resident CD8+ T have an increased need for FA metabolism for their function and survival^35.36^. We discovered that colon TRM CD8+ T cells depend on FAO for homeostasis in a similar manner. We showed that in PWH on ART, colon TRM CD8+ T cells exhibit a reduction in their intrinsic lipid reserves due to decreased PPARγ signaling, similar to other reports of the central role of PPARs in LD biogenesis and FAO^23–25^. To compensate for these diminished reserves, TRM CD8+ T cells scavenge lipids from the plasma membranes of nearby epithelial cells. This transfer of epithelial lipids contributes to T cell metabolic homeostasis via a PPAR-independent mechanism, but results in epithelial damage, leading to epithelial cell apoptosis and barrier compromise.

There are several limitations of this study. The specific mechanisms by which intestinal CD8+ T cells adapt to the metabolic demands during HIV infection or chronic inflammation merit further investigation. It has been recently described that tumor cells maintain their survival and proliferation in nutrient poor conditions by scavenging extracellular protein and lipids from adjacent cells^37–39^. Similarly, immune cells are known to feed off other cells through trogocytosis, the transfer of plasma membrane fragments from one cell to another^40^. Trogocytosis may be particularly important in the colon where CD8+ T cells are highly activated and maintain elevated states of metabolic demand^41^. Employing state-of-the-art heterotypic co-culture organoid assays, we characterized how colon resident CD8+ T cells acquire lipids from adjacent epithelial plasma membranes. However, the specific mechanistic details of this lipid scavenging, and the pathways of lipid utilization require further study. Additionally, although we demonstrated that treatment with rosiglitazone, an approved drug used for the management of diabetes and metabolic syndrome, can reverse the observed defects in TRM CD8+ T cells from PWH *ex vivo*, whether this would be effective in humans remains to be assessed.

We found that people with HIV (PWH) have persistent intestinal epithelial damage, evidenced by elevated plasma intestinal fatty acid binding protein (I-FABP) levels in both ART-naive and -treated individuals. Consistent with this, colonic biopsies showed increased epithelial apoptosis in PWH, despite unchanged intestinal stem cell frequencies. This epithelial cell apoptosis was recapitulated in a colon organoid model derived from biopsies from PWH and was dependent upon co-culture with autologous colon tissue resident memory (TRM) CD8+ T cells. These cells exhibited downregulation of PPARs and lipid metabolism pathways and impaired fatty acid oxidation (FAO). Treatment of colonoids co-cultured with autologous TRM CD8+ T cells with the PPARγ agonist rosiglitazone restored lipid homeostasis and ameliorated CD8+ T cell-induced epithelial apoptosis. Further, genetic ablation of PPARs in CD8+ T cells in a mouse model recapitulated the important role of these lipid regulators in maintaining epithelial homeostasis and gut barrier function. These findings may lead to the development of novel therapies to help improve intestinal barrier integrity and combat the associated comorbidities in PWH as well as other disease that impact intestinal homeostasis.

## METHODS

### Study design

We recruited 20 healthy, HIV-uninfected individuals and 20 people with HIV (PWH) on antiretroviral therapy (ART) for lower endoscopy (colonoscopy). This study (protocol number 2007p002102) was approved by the Massachusetts General Hospital (MGH) institutional review board, with informed consent obtained in writing after council with MGH clinical research coordinators and study physicians. The recruitment criteria for PWH on ART was those with plasma viral load below limit of detection (< 20 copies/mL) for >1 year prior to colonoscopy and were undergoing ART treatment for a median of 7 years. Healthy participants were recruited to closely match enrolled PWH based on age and sex. Individuals with a history of GI disease, clinical findings from a prior endoscopy, or who reported current or prior GI symptoms were strictly excluded.

### Sample collection

During colonoscopy, pinch biopsies were taken from the transverse colon and placed directly into collection media in the GI suite. Immediately prior to colonoscopy, blood was collected in ACD vacutainers (Beckton Dickinson) to isolate PBMCs and plasma. For Figure 1A, we used plasma from previously recruited participants and for Figure 1b and C, we used historically preserved FFPE biopsy pinches.

### Epithelial cell isolation

Pinch biopsies were collected in RPMI with 250μg/mL Zosyn (MGH Pharmacy), 2.5μg/mL amphotericin B (Sigma Aldrich), penicillin/streptomycin, 10mM HEPES, and L-glutamine and processed immediately. Pinches were washed with cold phosphate-buffered saline (PBS), and then incubated with 10mM ethylenediaminetetraacetic acid (EDTA Sigma Aldrich) for 30 minutes in the rotator in an 37C incubator.

### Methods for Organoid Culture

Organoids were prepared from crypts isolated from biopsy pinches as described before^42^. Isolated crypts were centrifuged at 100g for 3 min. The crypts were embedded in Matrigel (Corning) and incubated at 37° C for 20 min. The media formulation listed below was applied to the solidified Matrigel plugs. For subsequent passaging, the Matrigel plugs were mechanically dissociated by pipetting and exposed to TrypLE (Gibco) for 3 min, centrifuged and resuspended in Matrigel. Media for the organoids was based on L-WRN cell conditioned media (L-WRN CM). Briefly, L-WRN CM was generated by collecting 8 days of supernatant from L-WRN cells, grown in Advanced DMEM/F12 (Gibco) supplemented with 20% fetal bovine serum (Hyclone), 2 mM GlutaMAX, 100 U/mL of penicillin, 100 µg/mL of streptomycin, and 0.25 µg/mL amphotericin. L-WRN CM was diluted 1:1 in Advanced DMEM/F12 (Gibco) and supplemented with additional RSPO-1 conditioned media (10% v/v), generated using Cultrex HA-R-Spondin1-Fc 293T cells. Wnt activity of the conditioned media was assessed and normalized between batches via luciferase reporter activity of TCF/LEF activation (Enzo Leading Light Wnt reporter cell line). The following compounds were also added to the growth media: B27 (Gibco), 10 uM nicotinamide (Sigma-Aldrich), 50 ng/mL EGF (Novus Biologicals), 500 nM A83-01 (Cayman Chemical), 10 uM SB202190 (Cayman Chemical), and 500 nM PGE2 (Cayman Chemical). Y-27632 10uM (Cayman Chemical) was added to crypts only on their initial isolation.

### Immunofluorescence staining of organoids

Organoids were cultured on tissue culture-treated coverglass in 8-chambered slides (Ibidi). Organoids were washed in PBS, then fixed in 4% PFA (Electron Microscopy Sciences) in PBS for 30 minutes at room temperature. After fixation, cells were washed twice in PBS. Organoids were permeabilized for 15 minutes at room temperature using 0.1% Triton X-100 (Sigma Aldrich). Organoids were again washed twice with PBS. Protein blocking was done with 5% goat serum in PBS for one hour at room temperature. Serum was removed, and organoids were incubated with primary antibodies diluted in antibody diluent (Dako) overnight at 4°C. After primary antibody incubation, organoids were washed five times in PBS for five minutes each at room temperature. Organoids were then incubated in fluorophore conjugated secondary antibodies (1:300, ThermoFisher) diluted in 5% goat serum for 3 hours at room temp. Organoids were washed five times in PBS for five minutes each at room temperature. To visualize nuclei, organoids were stained with NucBlue fixed cell reagent (ThermoFisher) for 10 minutes at room temperature. Organoids were washed one time in PBS, and then imaged on a Zeiss Cell Discoverer LSM900 confocal microscope.

### LDH assay for organoid culture

LDH secretion was measured from organoid culture media according to manufacturer’s protocol.

### ELISA

The concentration of some of the plasma markers was measured by ELISA using Human FABP2/I-FABP DuoSet ELISA (R&D Systems, DY3078), per manufacturer’s instructions.

### PBMC isolation

Peripheral blood mononuclear cells (PBMCs) were isolated from the blood of HIV-uninfected and PWH by density gradient centrifugation. Blood was transferred to a 50 mL conical and centrifuged at 2600 rpm for 15 minutes. The resultant plasma layer was discarded, and the remainder was volumed up to 30mL with Hank’s Balanced Salt Solution (HBSS). This mixture was then layered with 15mL of Histopaque-1077, after which the conical was spun at 1500 rpm for 45 minutes. The PBMC layer was then collected in a new tube, washed twice with HBSS, counted, and frozen in a 10% DMSO solution.

### Colonic immune cell isolation

Lymphocytes were isolated from intestinal biopsy pinches by modifying the protocol as described^48–50^. Biopsy pinches were transferred to a 50mL conical containing intraepithelial lymphocytes (IEL) stripping buffer (PBS, 10mM DTT, 5mM EDTA, 10mM HEPES, & 5% FCS) and incubated under continuous rotation for 20 minutes at 37°C. After separating tissue pieces from the supernatant, the tissue was again treated with IEL buffer and incubated while the supernatant was spun down and IELs were collected. This process was repeated until all IELs had been isolated from the digested biopsy tissue. Following IEL isolation, biopsy pinches were placed into a Collagenase digestion solution. Tissue pinches were incubated in Collagenase at 37°C for one hour to release lamina propria (LP) cells. These cells, in combination with isolated IELs, were enriched for CD45+ cells by positive immunomagnetic selection using Miltenyi CD45 microbeads. Cells were cultured in complete RPMI media (10% FBS) in the presence of 50 IU/mL human IL-2, 10 ng/mL human IL-7, and 10 ng/mL human IL-15. After 48 hours, cells were harvested for co-culture with organoids or processed by flow cytometry.

### Organoid-immune coculture

Immune cells and organoids were mixed in 100000 immune cells to 20 organoids ratio and embedded in matrigel for their co-culture.

### Flow cytometry

Cryopreserved PBMCs and colon derived immune cells were thawed, washed, and counted. 100,000 cells per sample were allotted into FACS tubes, along with 20,000 cells for each fluorescence minus one (FMO) control and mitochondrial dye compensation control tube. The following fluorescence-tagged antibodies were used for PBMCs: CD3-PerCP-Cy5.5, CD8-V500, CD45 RO-APC and LIVE/DEAD Fixable Blue Stain. Colonic immune cells were stained with the fluorescence-tagged antibodies: CD3-PerCP-Cy5.5, CD4-BUV395, CD8-V500, CD103-PECy7, S1PR1-PE, CD45RO-APC, CD69-FITC and LIVE/DEAD Fixable Blue Stain. All cells were incubated with Fc block for 5 minutes at room temperature (RT) and stained with surface antibodies for 20 minutes at RT. After washing, cells were processed by the flow cytometer (4 Laser LSR II). Compensation controls for each surface antibody were prepared using anti-mouse IgG compensation beads. Analysis of flow cytometry data was performed using FlowJo software. Fluorescence minus one controls were used to inform gating.

### Intracellular staining

Immune cells were in complete RPMI media for 6h in the presence of, anti-CD107a, 1μM GolgiStopTM (BD Biosciences), brefeldin A (5μg/mL, Sigma Aldrich). Cells were incubated for 10 minutes in PBS with 0.5mM EDTA prior to staining for surface markers and viability (Blue Viability dye, Invitrogen) for 20 minutes at room temperature. Cells were fixed in 4% paraformaldehyde (PFA) and permeabilized using FACS Perm 2 (BD Biosciences) prior to intracellular staining for IFN-γ, TNF, perforin and Granzyme B. After staining, cells were re-suspended in 1% PFA and stored at 4°C in the dark until analysis within 24h.

### Immunofluorescence staining of pinch biopsies

Biopsies were immediately preserved in formalin in the endoscopy suite. Pinches were transferred to 70% ethanol after 24 hours and embedded in paraffin blocks. 4μm sections were used for analysis. Sections were de-paraffinized using Histoclear (National Diagnostics) and graded alcohols. Heat-induced epitope retrieval was performed in DIVA decloaker (Biocare Medical) at 125°C for 30 seconds using a pressure cooker (Biocare Medical). Sections were stained with the antibodies overnight at 4°C. Slides were washed in TBS with 0.1% Tween-20 (TBST), and then incubated with AlexaFluor-conjugated secondary antibodies (ThermoFisher) for 2 hours at room temperature. Slide were washed with TBST, stained with NucBlue Fixed Cell Reagent (ThermoFisher), and mounted with ProLong Gold Antifade (ThermoFisher). Slides were imaged using the TissueFAXS whole-slide scanning system (Tissue Gnostics).

### Fatty acid oxidation assay

Flow cytometer sorted or magnetically enriched CD8+ T cells from human colon were plated on a 24 well plate in 400 μL RPMI with 5mM glucose and allowed to incubate in a humidified 37°C incubator at 5% CO_2_ for overnight. 10% essentially fatty acid free BSA (Sigma Aldrich, A6003) in PBS was complexed at a volume ratio of 6.7:3 with palmitic acid-[9,10-^3^H] (Pekin Elmer, NET043001MC) by vortexing for 60 s and was added at a 1:100 ratio to crypt medium. This hot medium was split into half and Etomoxir (4 μM final) was added to inhibit FAO. 100 μl of the hot FFA:PA:medium mixture was added to the incubated cells, for a total volume of 500 μl, and were incubated for 1 hour, 37°C, 5% CO_2_. Samples were removed from the wells and pelleted at 21K r*cf.* for 2:30 min. 400 μL of resulting supernatant was transferred to a filter column (Fisher Scientific, 11-387-50) containing 3 mL of activated Dowex®1X8 resin (Sigma Aldrich, 217425). 2.5 mL of ddH_2_O was added to elute ^3^H-water from the column. 750 μL of eluent was added to 2.5 mL of EcoLume (MP Biomedicals, 882470). Beta-counts were measured on a scintillation counter (Beckman Coulter, LS6500).

### Bulk sequencing of CD8+ T cells

#### RNA isolation, library construction, sequencing, and alignment

CD8+ T cells from PBMCs and colon were FACS sorted directly into 50 μL of RLT Lysis Buffer (QIAGEN) supplemented with 1% v/v 2-mercaptoethanol. Briefly, 50 μL of mixed lysate from each sample was transferred to a skirted 96 well plate. Genetic material was pulled down and purified by mixing the lysate in each well with 2.2x volumes of Agencourt RNAClean XP SPRI beads (Beckman Coulter) and washing 3x with 75 μL of 80% ethanol. After drying, the SPRI beads were re-suspended in 4 μL of pre-reverse transcription (RT) mix, incubated for 3 min at 72°C, and placed on ice. Next, Smart-Seq2 Whole Transcriptome Amplification (WTA) was performed: 7 μL of RT mix was added to each well and RT was carried out; then, 14 μL of PCR mix was added to each well and PCR was performed. Thereafter a cDNA cleanup was performed using 0.6x and 0.8x volumes of Agencourt AMPure XP SPRI beads (Beckman Coulter) which was then quantified using a Qubit dsDNA HS Assay Kit (Life Technologies). Library size and quality were measured by Bioanalyzer using a High Sensitivity DNA Analysis Kit (Agilent Technologies). Sequencing libraries were prepared from WTA product using Nextera XT (Illumina). After library construction, a final AMPure XP SPRI clean-up (0.8 volumes) was conducted. Library concentration and size were measured with the KAPA Library Quantification kit (KAPA Biosystems) and a TapeStation (Agilent Technologies), respectively. Finally, samples were sequenced on a NextSeq500 (30 bp paired end reads) to an average depth of 5 million reads. Reads were aligned to hg38 (Gencode v21) using RSEM and TopHat^51^ and estimated counts and transcripts per million (TPM) matrices generated. Any samples with fewer than 5×10^5^ or more than 6×10^6^ aligned reads or fewer than 10,000 uniquely expressed genes were removed from subsequent analysis.

#### RNA-Seq Differential Expression Analysis

Differential expression analysis was performed using DESeq2 (v1.18.1)^52^. Expected counts from biological replicates for each cell type and participant were averaged prior to differential expression to prevent participant specific genes from generating false positives and reduce spurious heterogeneity from small (100-cell) populations. Small populations may show skewed expression based on the cell composition within; thus, this replicate averaging approach is particularly important given our limited access to pediatric tissue sources and low frequency of these immune populations in order to remove further bias from small population sorts.

Gene set analysis was performed using Ingenuity Pathway Analysis (IPA; Winter 2019 Release, QIAGEN Inc.) and Gene Set Enrichment Analysis (GSEA) using the piano package in R (1.18.1). For IPA, DEGs whose FDR corrected q < 0.1 were used in the “Core” analysis with the log2FC and q values included in the analysis. To implement GSEA on our DESeq2 results, we used the log2FC of all genes whose FDR corrected q < 0.1 as t-value input into the runGSA function with setting the argument geneSetStat = “gsea.” We chose to use the KEGG and GO databases (downloaded from MSigDB v7.0)53 for GSEA analysis as these databases are well annotated for metabolic and cellular activation gene sets that are cell-type agnostic.

### Seahorse mitochondrial metabolic analysis

#### Cell preparation and plating

The Agilent Seahorse XFp Cell Mito Stress Test was performed according to manufacturer’s protocol using the Seahorse XFp Analyzer. Peripheral and GI mucosal CD8+ T cells were purified from PBMCs and GI mucosal lymphocytes, respectively, by positive magnetic cell separation. CD8+ T cells were washed in XF medium, counted, and resuspended in XF medium at a concentration of 5×10^6^ cells/mL. 2×10^5^ cells/well were plated in a 96-well tissue culture plate, centrifuged at 1500rpm for 5 minutes, and covered with 140 μL of XF medium. The plate was placed in a non-CO2 incubator overnight at 37 degrees Celsius.

#### Cartridge Preparation

200μL of calibrant was added to each well of the Seahorse XF Cell Culture Miniplate. The miniplate was coated with Poly-D-lysine and incubated overnight at 37°C in non-CO2 conditions. Cells were transferred to the Miniplate the following day.

#### Program Preparation

On day 2, the Mito Stress test was programmed in the Seahorse console:

1. Calibrate
2. Equilibrate
3. Base Line Readings (3x Loop)
4. Mix 1 → 3 min Wait → 2 min Measure → 3 min End
5. Inject Port B (3x Loop)
6. Mix 2 → 3 min Wait → 2 min Measure → 3 min End
7. Inject Port C (3x Loop)
8. Mix 3 → 3 min Wait → 2 min Measure → 3 min End
9. End

Reagents were prepared at the following concentrations: 1μM oligomycin, 1.5 μM FCCP, 100nM rotenone, and 1μM antimycin A. Drugs were then loaded into the delivery ports of the sensor cartridge with a multichannel pipette:

Port B: 20μL oligomycin
Port C: 22μL FCCP
Port D: 24μL rotenone + antimycin A

### Liposomal nanoparticles

Liposomes composed of 1,2-dioleoyl-sn-glycero-3-phosphocholine (DOPC, Avanti #850375), cholesterol (Avanti #700100), and 1,2-dioleoyl-sn-glycero-3-phosphoethanolamine-N-(Cyanine 5) (18:1 Cy5 PE, Avanti #810335) in a 51.5:38.5:10 mole ratio were synthesized by lipid film rehydration in phosphate buffered saline. The liposomes underwent six freeze/thaw cycles using liquid nitrogen to ensure unilamellarity and were subsequently extruded through a 100nm and 200nm nm membrane to ensure even size distribution, which was confirmed via dynamic light scattering.

### Mice

Mice were under the husbandry care of the Department of Comparative Medicine in the Koch Institute for Integrative Cancer Research. All procedures were conducted in accordance with the American Association for Accreditation of Laboratory Animal Care and approved by MIT’s Committee on Animal Care. The following strains were obtained from the Jackson Laboratory: Ppardfl/fl (B6.129S4-Ppardtm1Rev/J, stock number 005897), Ppargfl/fl ( B6.129-Ppargtm2Rev/J, stock number 004584), Ppara−/− (129S4/SvJae-Pparatm1Gonz/J, stock number 003580) and Cd8a-cre (C57BL/6-Tg(Cd8a-cre)1Itan/J, stock number 008766). All mice were sex and aged matched and provided food *ad libitum*.

### Crypt Isolation from mice colon and organoid culture

Colon crypts were isolated from wild type C57BL/6 mice as described previously^43^. Briefly, colons were washed with phosphate-buffered saline (PBS), minced into approximately 1cm segments, and incubated in 8mM EDTA for 45 minutes at 37°C. After incubation, EDTA was replaced with PBS and epithelial crypt were obtained by vigorous shaking. The cell suspension was passed through a 70μm filter. Crypts were resuspended in the appropriate volume of one-third primary culture media and two-thirds Matrigel (growth factor reduced, phenol red-free, Corning). Ten microliter droplets of crypts were plated in 48-well tissue culture treated plates (Genesee Scientific). Matrigel was allowed to solidify for 30 minutes at 37°C, after which 0.5mL of primary culture media was applied. Primary organoids (crypt-derived) were cultured in advanced DMEM/F12 (Gibco) supplemented with 5% fetal calf serum (FCS), Antibiotic/Antimycotic (Gibco), GlutaMax (ThermoFisher), 1X B-27 (ThermoFisher), 50ng/mL EGF (Peprotech), 10μM Y-27632 (StemCell Technologies), 1% Noggin-conditioned medium, and 2% R-spondin1-conditioned medium (“primary culture medium”). Organoids were cultured for at least 3 days at 37°C in a humidified, sterile incubator with 5% CO_2_.

### Statistical Analysis

Data points represent biological replicates for mouse samples and individual participants in case of human samples and are shown as the means +/- SD. Statistical significance was determined as indicated in the figure caption using two-tailed, unpaired or paired Student’s t tests or one-or two-way analysis of variance (ANOVA) (for greater than two comparison groups). Statistical details of each experiment can be found in the figure captions. P values of <0.05 were considered significant and represented by asterisk as follows: \**P* ≤ 0.05, \*\**P* ≤ 0.01, \*\*\**P* ≤ 0.001, \*\*\*\**P <* 0.0001. Analyses were performed using Prism 9.0 (Graph-Pad software).

## Acknowledgements

We thank the study participants for donation of clinical samples, Daniel Worrall and the Ragon Institute Clinical Team for clinical study support, the study participants and Michael Waring for flow cytometry assistance.

## Author Contributions

Conceptualization: U.D.A., O.H.Y., A.E.R., and D.S.K.; investigation: U.D.A., L.M.F., A.N.P., H.B., P.S., A.H.; generation of liposomal nanoparticles: B.K., B.R. D.J.I.; microscopic imaging assistance: T.J.D.; electron microscopy: M.L., P.B.; RNA seq analysis: Q.Z., S.N., A.S.; procurement of clinical specimens: O.A.,F.M., S.K., H.K.; writing (original draft): U.D.A.; review and editing: D.S.K and O.H.Y and other authors

