## Supplementary figures and legends for "PPARγ downregulation in colonic CD8+ T cells results in epithelial barrier disruption in people with HIV on antiretroviral therapy"

Supplementary 1

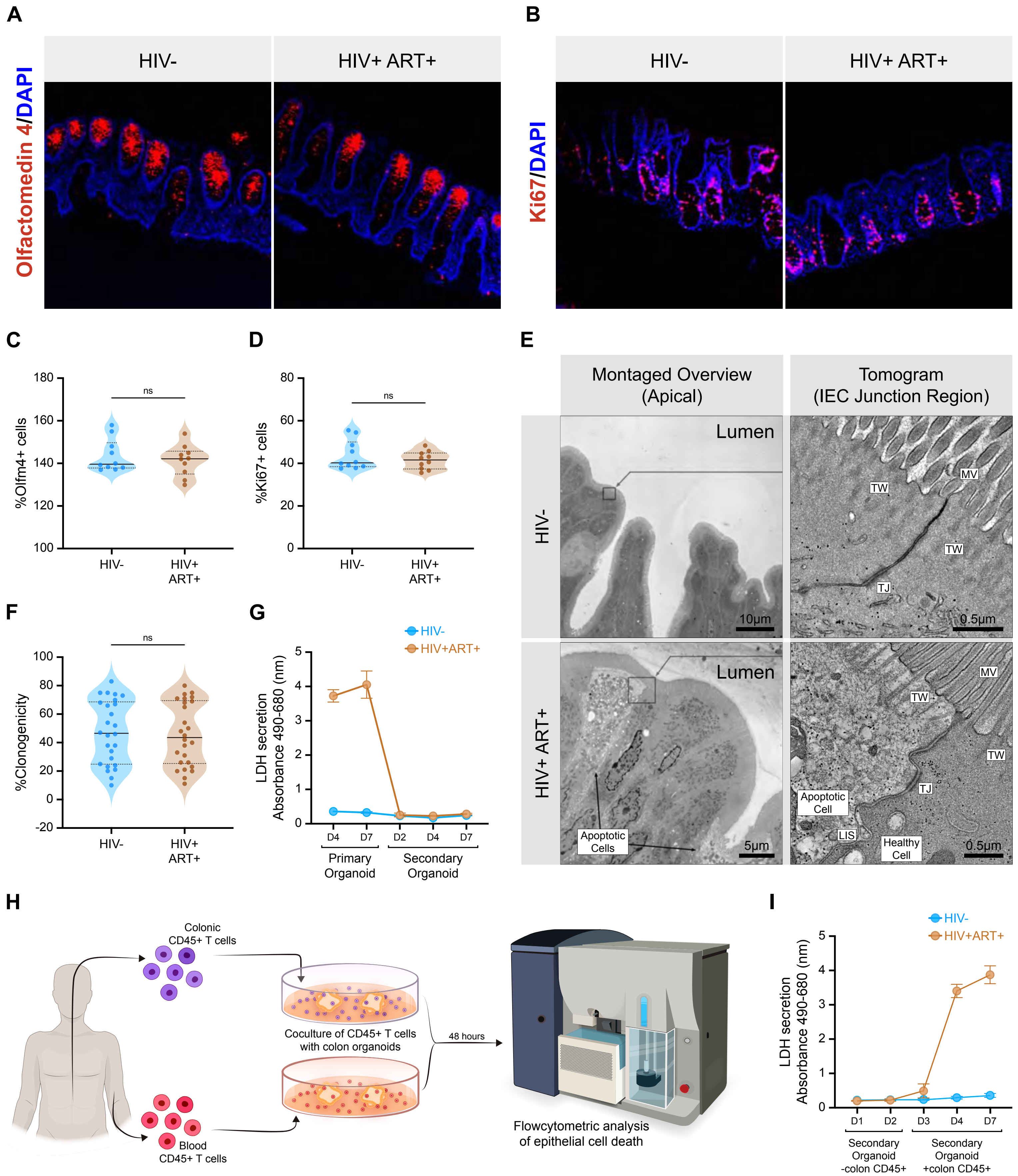

Supplementary 2

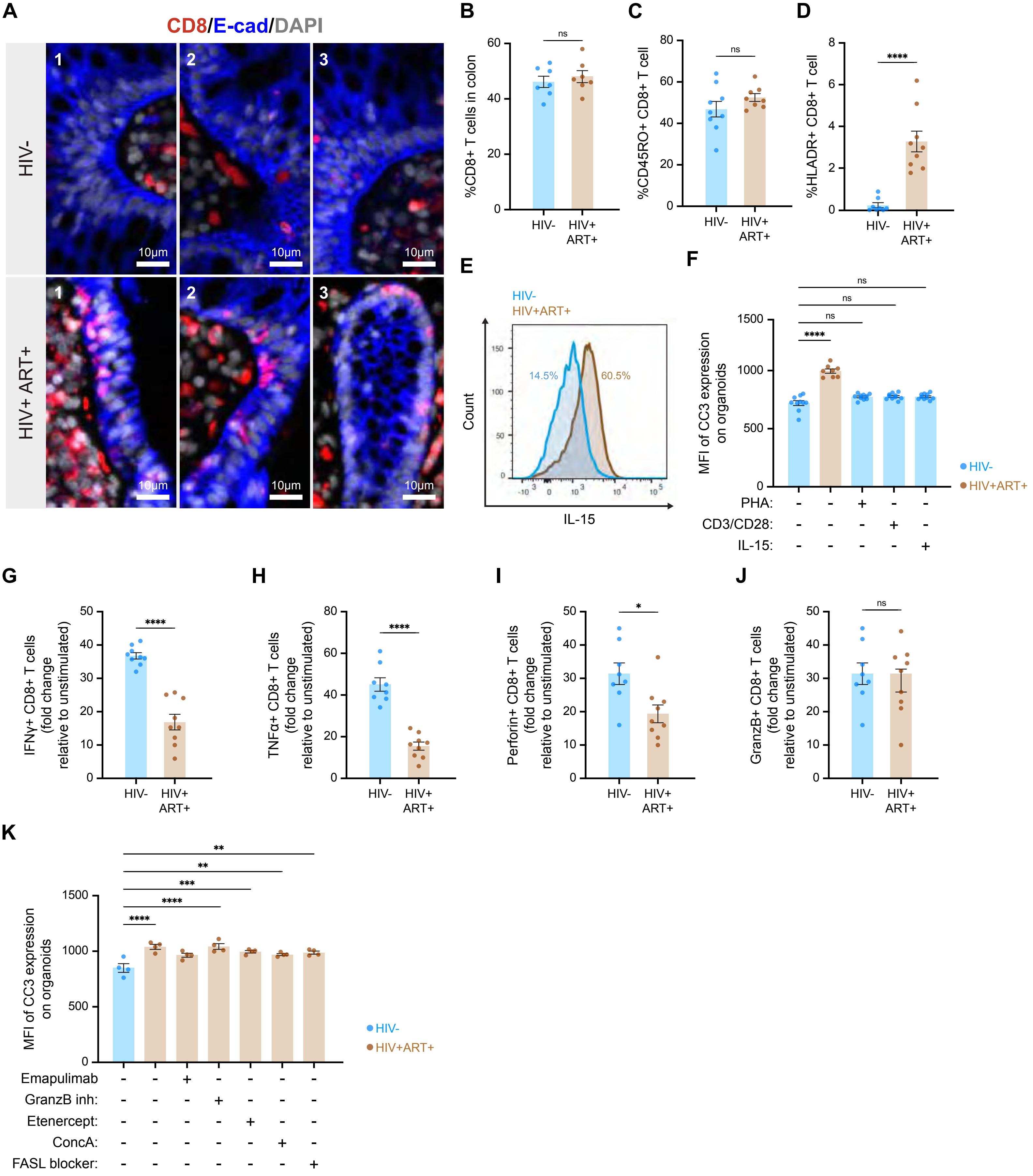

Supplementary 3

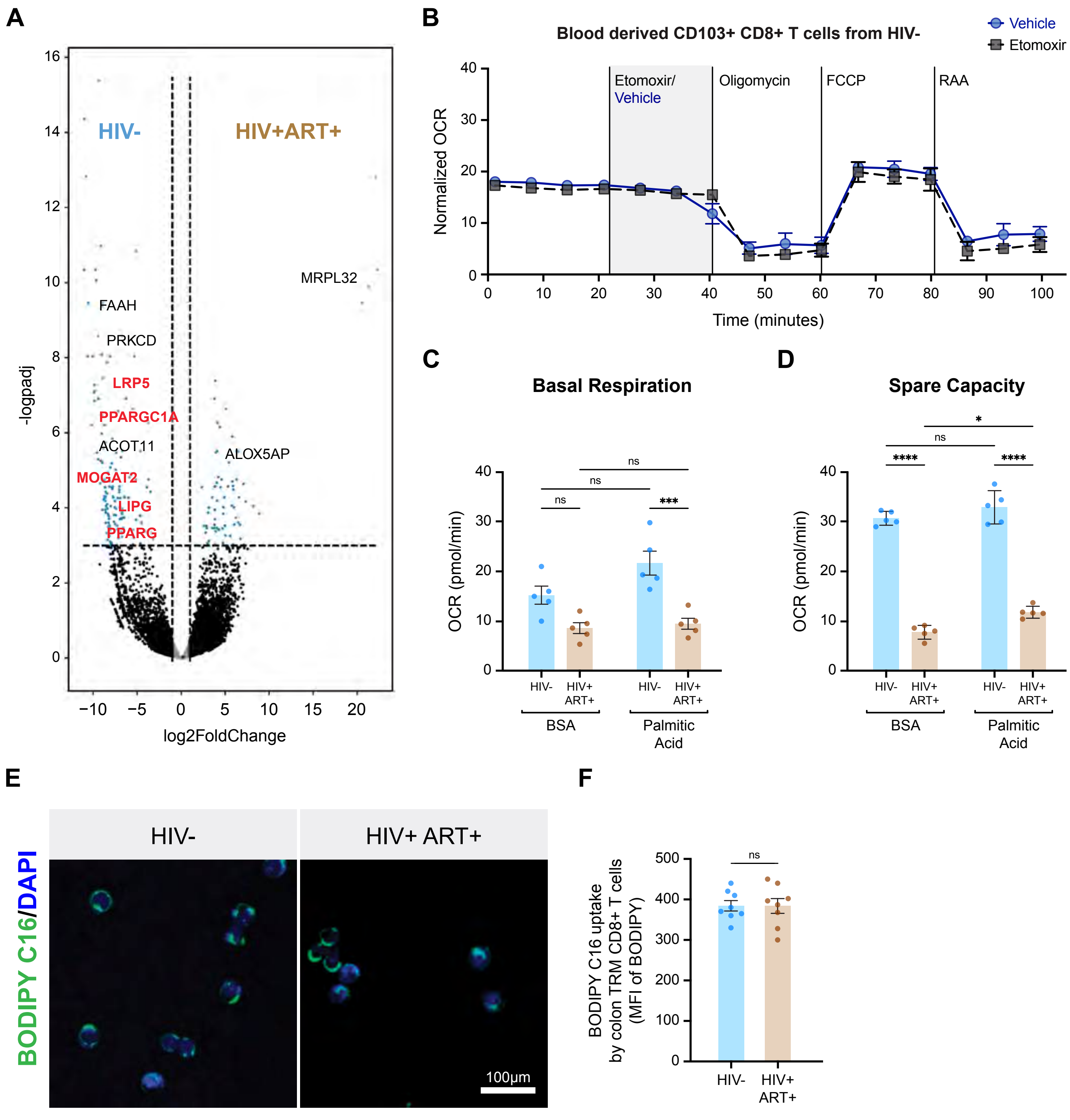

Supplementary 4

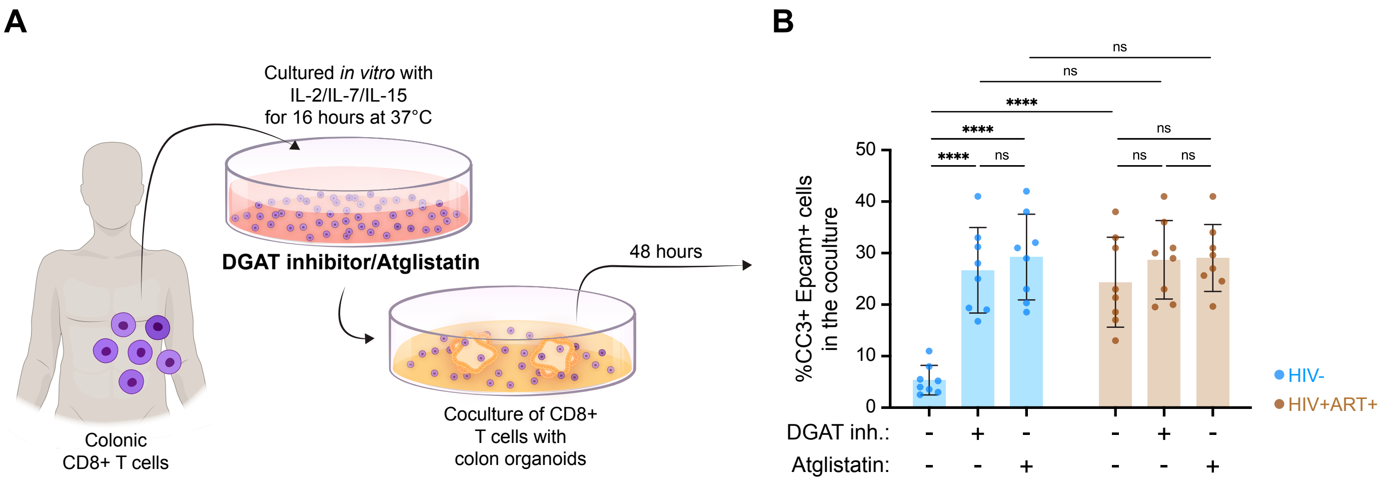

Supplementary 5

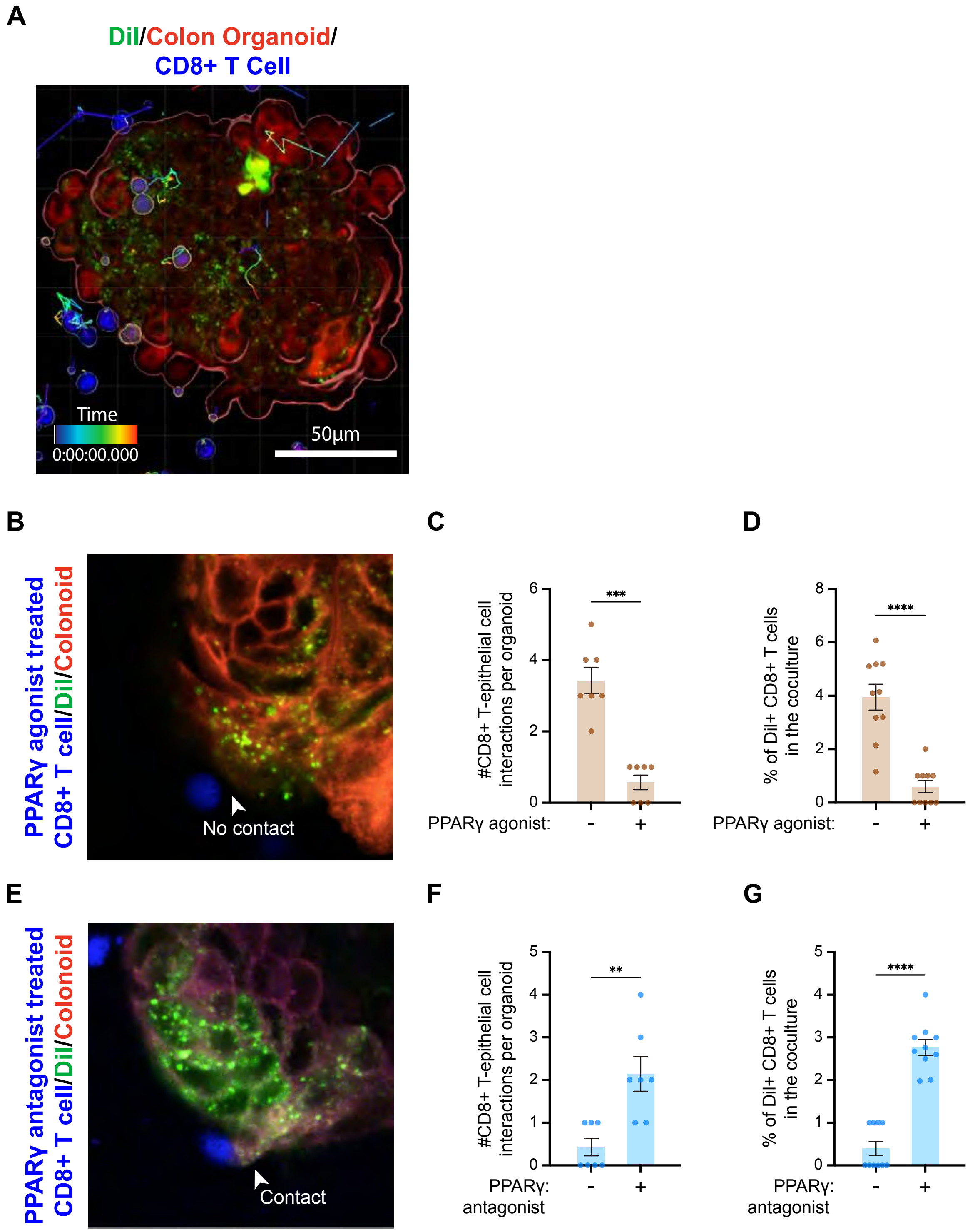

Supplementary 6

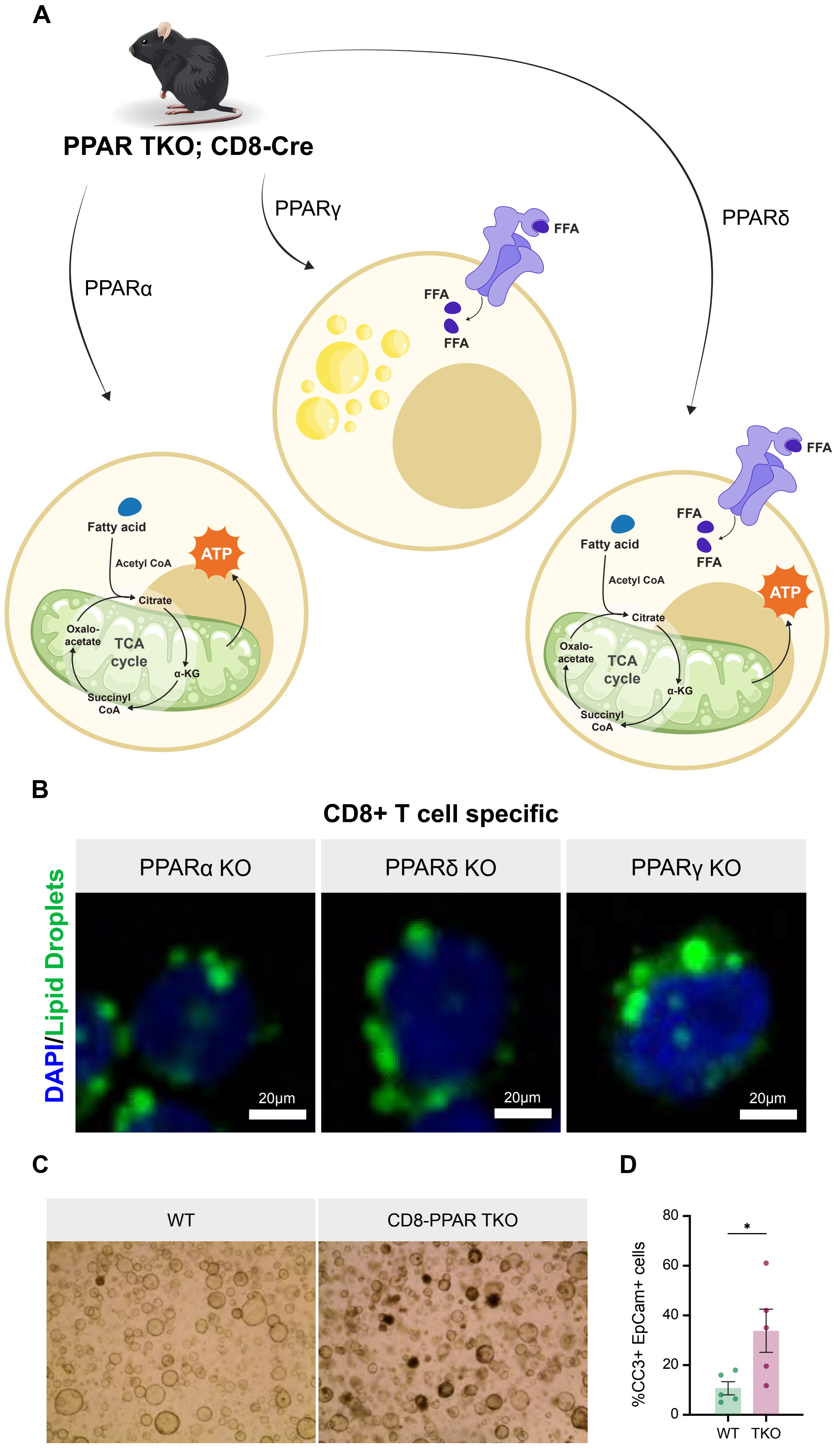

**Supplementary Figure 1**

A. Confocal microscopic images of colonic biopsy showing epithelial progenitor cells (Olfactomedin4 in red) of PWH with ART and uninfected individuals. Scale bars 50μm
B. Confocal microscopic images of colonic biopsy showing proliferative cells in the crypts (Ki67 in red) of PWH with ART and uninfected individuals. Scale bars 50μm
C. Quantification of confocal imaging for colonic epithelial progenitor cells (Olfm4+) in PWH with ART and uninfected individuals.
D. Quantification of confocal imaging for colonic epithelial proliferative cells (Ki67+) in PWH with ART and uninfected individuals.
E. Electron Microscopy showing intact tight junctions in apoptotic epithelium of ART treated PWH compared to uninfected
F. Quantification of clonogenic potential of colonic epithelial cells from primary colonoids of PWH with ART and uninfected individuals.
G. Spontaneous LDH secretion of primary vs secondary colonoids from HIV uninfected and PWH.
H. Schematic depicting the isolation of autologous blood and colon-derived immune cells from patients, followed by coculture with the patient's corresponding autologous secondary colonoids
I. Spontaneous LDH secretion of secondary colonoids from HIV uninfected and PWH after they were cocultured with corresponding patient autologous blood and colon derived immune cells.

**Supplementary Figure 2**

A. Confocal microscopy images of colon biopsy pinches showing the localization of CD8+ T cells and the epithelial region; Insets 10$\mu$m
B. Quantification of CD8+ T cells isolated from colons of PWH with ART and uninfected individuals.
C. Quantification memory population of CD8+ T cells (CD8+ CD45RO+) isolated from colons of PWH with ART and uninfected individuals.
D. Quantification of frequency late activated colon TRM CD8+ T cells (CD8+ CD45RO+ CD103+ HLADR+) from PWH with ART and uninfected individuals.
E. Histogram showing intracellular IL15 expression between colon derived immune cells from PWH on ART (brown) and HIV-uninfected (blue).
F. Quantification of MFI (mean florescence intensity) of CC3 expression on the organoids that are cocultured with colon TRM CD8+ T cells derived from healthy, HIV-uninfected (blue bar), PWH on ART (brown bar), healthy, HIV-uninfected and treated with PHA, CD3/CD28 and IL15 respectively (blue bars).
G. Quantification of frequency of IFNg+ colon TRM CD8+ T cells in HIV uninfected healthy donors and PWH on ART
H. Quantification of frequency of TNFa+ colon TRM CD8+ T cells in HIV uninfected healthy donors and PWH on ART
I. Quantification of frequency of Perforin+ colon TRM CD8+ T cells in HIV uninfected healthy donors and PWH on ART
J. Quantification of frequency of GranzymeB+ colon TRM CD8+ T cells in HIV uninfected healthy donors and PWH on ART

K. Quantification of MFI (mean florescence intensity) of CC3 expression on the organoids that are cocultured with colon TRM CD8+ T cells derived from healthy, HIV-uninfected (blue bar), PWH on ART (brown bar), PWH on ART treated with mAb for IFNg (Emapulimab), GranzymeB inhibitor, TNF inhibitor (Etanercept), perforin inhibitor (ConcanamycinA) and FASL blocker respectively (brown bars).

**Supplementary Figure 3**

A. Volcano plot showing downregulation of lipid metabolism associated genes in PWH on ART (n= 8) compared to uninfected individuals (n=4) from Durban, South Africa patient cohort.
B. Oxygen consumption rate (pmol/min/cells) of sorted blood derived CD103+ CD8+ T cells from healthy, HIV-uninfected individuals was performed by SeaHorse assay to evaluate whether they perform FAO. The OCR was measured during the mitochondrial stress test with the addition of oligomycin, carbonyl cyanide p-trifluoromethoxy phenylhydrazone (FCCP), and rotenone and antimycin A drugs (RAA). Results were normalized with the cell count.
C. Basal OCR was calculated.
D. Spare capacity was calculated.
E. Confocal images of colon TRM CD8+ T cells depicting fatty acid uptake capacity by internalizing BODIPY C16 when cultured with BODIPY C16.
F. Quantification of MFI of BODIPY uptake by colon TRM CD8+ T cells.

**Supplementary Figure 4**

A. Schematic showing colon TRM CD8+ T cell treated with DGAT inhibitor and ATGL inhibitor (Atglistatin) then cocultured with autologous patient derived secondary colonoids.

B. Flowcytometric quantification of epithelial apoptosis (Epcam+/ CC3+) in coculture, when colon TRM CD8+ T cell of PWH on ART and uninfected individuals was supplemented with DGAT inhibitor and ATGL inhibitor (Atglistatin).

**Supplementary Figure 5**

A. Diagram showing epithelial cells stained with Cell Mask (red, outlining the organoid) and their lipids stained with DiI (green), cocultured with colon TRM CD8+ T cells stained with Cell Trace Violet (blue) for live imaging using a confocal microscope.

**B.** Snapshot of live imaging of colon TRM CD8+ T cells cocultured with colonoids showed: Colon TRM CD8+ T cells (blue) derived from PWH on ART, when treated with PPARγ agonist showed no interaction with epithelial cell (red) of colonoid.

**C.** Quantification of the frequency of colon TRM CD8+ T cell interacting epithelial cells in the coculture of PPARg agonist treated colon TRM CD8+ T cell and patient autologous secondary colonoids derived from PWH on ART.

**D.** Quantification of the frequency of DiI+ colon TRM CD8+ T cell in the coculture of PPARγ treated colon TRM CD8+ T cell and patient autologous secondary colonoids derived from PWH on ART.

**E.** Snapshot of live imaging of colon TRM CD8+ T cells cocultured with colonoids showed: Colon TRM CD8+ T cells (blue) derived from uninfected individuals, when treated with PPARγ antagonist showed interaction with epithelial cell (red) of colonoid.

**F.** Quantification of the frequency of colon TRM CD8+ T cell interacting epithelial cells in the coculture of PPARg antagonist treated colon TRM CD8+ T cell and patient autologous secondary colonoids derived from uninfected individuals.

**G.** Quantification of the frequency of DiI+ colon TRM CD8+ T cell in the coculture of PPARγ antagonist treated colon TRM CD8+ T cell and patient autologous secondary colonoids derived from uninfected individuals.

**Supplementary Figure 6**

A. Schematic showing the CD8+ T cell specific knock out for PPARα, δ, γ and α/δ/γ TKO

B. Confocal imaging of lipid droplets (green) in colonic CD8+ T cells from WT and CD8+ T cell specific PPARα, δ, γ KO mice. Scale bar 20mm

C. Bright Field images of WT mouse colonoids when cocultured with colonic CD8+ T cells from WT and CD8+ T cell specific TKO mice. The cell death in the organoids is visualized as dark spots.

D. Quantification of flow cytometric analysis of colonic epithelial apoptosis (CC3+ Epcam+) from the colonoid cultures.
